# Modulating Peripheral Neural Activity: Prolonged Low-Intensity Ultrasound for Controlled Excitation and Suppression in Rat Sciatic Nerve

**DOI:** 10.1101/2024.12.26.630403

**Authors:** Heba M. Badawe, Pierre D. Mourad, Massoud L. Khraiche

## Abstract

**Objective:** Low-intensity, low-frequency ultrasound has shown promise for neuromodulation, particularly for influencing peripheral neural activity. However, the precise parameters required to modulate neuronal activity consistently remain poorly understood, limiting its broader application. Here, we investigate the effects of varying sonication duration (SD) and duty cycle (DC) on motor neuronal responses in the rat sciatic nerve, focusing on understanding how cumulative energy exposure influences the activation, enhancement, or suppression of peripheral neural activity during ultrasound neuromodulation.

**Approach:** We apply low-intensity, low-frequency ultrasound to the rat sciatic nerve in vivo at different sonication durations (30s, 60s, 90s, and 120s) and duty cycles (30%, 50%, and 80%). The cumulative energy exposure is calculated as the product of spatial-peak pulse-average intensity, SD, and DC. Electromyographic (EMG) activity in the gastrocnemius muscle is measured, and the thermal effects are monitored to ensure a non-cavitational, non-thermal application.

**Main Results:** Our findings demonstrate that higher cumulative energy exposures suppress EMG activity in the gastrocnemius muscle (enervated by the sciatic nerve). However, lower cumulative energy exposures enhance EMG activity and motor stimulation. Notably, the ultrasound-induced EMG changes persisted for 5 minutes post-sonication – three to five times longer than the application duration -- underscoring the therapeutic potential of ultrasound for precise neural control. In vivo evaluations suggest the mechanical nature of the observed effects without any significant temperature increase or induction of cavitation. In vivo evaluations suggest the mechanical nature of the observed effects without any significant temperature increase or induction of cavitation.

**Significance:** Interestingly, our results show a switch from excitation to suppression of electrically evoked EMG activity following ultrasound sonication depending on the acquired cumulative energy. This study establishes a safe parameter space for prolonged neuromodulation, demonstrating its potential for therapeutic applications that can precisely modulate peripheral nervous system activity. These findings contribute to the development of ultrasound-based treatments for neurological conditions, offering a novel and controllable method for peripheral nerve stimulation.

## 1. Introduction

Acoustic neuromodulation began with foundational studies in the 1950s and 1960s, when researchers like Fry et al. discovered that focused ultrasound could selectively affect neural tissues, initially exploring its potential for brain lesioning [1]. However, it was not until technological advances in the 2000s, particularly in low-intensity focused ultrasound (LIFU), that non-invasive neuromodulation became feasible. Renewed studies demonstrated that LIFU could modulate neuronal activity without causing damage, leading to expanded applications in treating neurological and psychiatric conditions. In the 2010s, human studies further validated ultrasound’s potential in conditions such as essential tremor, Parkinson’s disease, and depression [2, 3].

Today, ultrasound neuromodulation has emerged as a promising non-invasive technique for modulating neuronal activity in both the central and peripheral nervous systems. As an alternative to electrical and pharmacological neuromodulation methods, ultrasound offers a high degree of spatial precision with minimal invasiveness, making it an attractive option for therapeutic applications. Despite these advantages, a comprehensive understanding of the specific ultrasound parameters required to achieve consistent excitatory or inhibitory effects remains elusive, particularly in the peripheral nervous system (PNS). Previous studies have shown varied results, with some reporting enhanced nerve activity Today, ultrasound neuromodulation has emerged as a promising non-invasive technique for modulating neuronal activity in both the central and peripheral nervous systems. As an alternative to electrical and pharmacological neuromodulation methods, ultrasound offers a high degree of spatial precision with minimal invasiveness, making it an attractive option for therapeutic applications. Despite these advantages, a comprehensive understanding of the specific ultrasound parameters required to achieve consistent excitatory or inhibitory effects remains elusive, particularly in the peripheral nervous system (PNS). Previous studies have shown varied results, with some reporting enhanced nerve activity [4–10] and others suppression [11–14] or no effect [15]. Add to that, human studies of focused ultrasound applied to peripheral nerves demonstrate its ability to induce. Add to that, human studies of focused ultrasound applied to peripheral nerves demonstrate its ability to induce [16, 17], or suppress [18] sensations, as well as to simultaneously induce both sensations and motor activity [19]. This highlights a critical gap in standardized methodologies for ultrasound neuromodulation.

This study addresses this gap by exploring the effects of cumulative energy exposure, defined as a function of sonication duration (SD) and duty cycle (DC), on the modulation of electrically evoked motor neuronal responses in the rat sciatic nerve. By investigating both thermal and non-thermal mechanisms in a controlled, non-cavitational environment, this work aims to establish a parameter space for safe and sustained PNS modulation. Our findings suggest a potential paradigm for ultrasound-induced neuromodulation, where tailored energy exposure parameters can yield precise excitatory or inhibitory outcomes without thermal damage. These insights contribute to the development of ultrasound-based therapies for managing neurological and peripheral nerve disorders, presenting a non-invasive, controllable method for peripheral nerve stimulation with significant clinical implications.

## 2. Materials and Methods

### 2.1. Animal Preparation and Surgical Procedure

All experimental procedures onon animals were approved by the Institutional Animal Care and Use Committee at the American University of Beirut, Lebanon. Twenty male Sprague-Dawley rats, weighing between 270-480 g, were used in this study. The rats were kept under a 12-hour day/night cycle at a temperature of 25°C. Anesthesia was induced using a combination of 80 mg/kg Ketamine and 20 mg/kg Xylazine, administered intraperitoneally. Following anesthesia, the fur over the surgical site was shaved, and the left sciatic nerve was exposed by making an incision through the shaved skin. The sciatic nerve was identified lateral to the femur bone, just below the biceps femoris muscle. Once isolated from the surrounding tissues in a “hammocked” way, excess skin and muscles were retracted using skin flaps. Saline was added to create a pool under the sciatic nerve to prevent desiccation. The depth of anesthesia was monitored by periodically testing the rat’s pedal reflex through frequent pinching of the forelimb pad. Any necessary redosing of anesthesia was administered before initiating the experiment.

### 2.2. Electrode Placement and Neurophysiological Recordings Setup

A bipolar, hooked, Teflon-coated, silver-stimulating electrode was coiled around the “hammocked” nerve for electrical stimulation. A needle, uncoated stainless steel, recording electrode was inserted in the gastrocnemius muscle to record EMG activity in response to the induced electrical stimulation. A reference electrode was affixed to the excess skin at the back of the rat for grounding. The ultrasound-emitting transducer, once coupled with a cone filled with ultrasound gel, was precisely positioned above the exposed nerve, and placed between the stimulating and recording electrodes (Figure 1A). The stimulating electrode received square electrical signals of 2 Hz frequency, a pulse duration of 2 ms, and a pulse amplitude of 1 V from a single-channel GRASS S44 Stimulator coupled with a GRASS SIU5 stimulus isolation unit. EMG signals were recorded during electrical and ultrasound stimulation of the gastrocnemius muscle through the recording electrode connected to an isolated bio-amplifier (WPI) of 10^3^ gain and band-pass filtered from 10 Hz to 3 kHz (Figure 1B).

**Figure 1.**
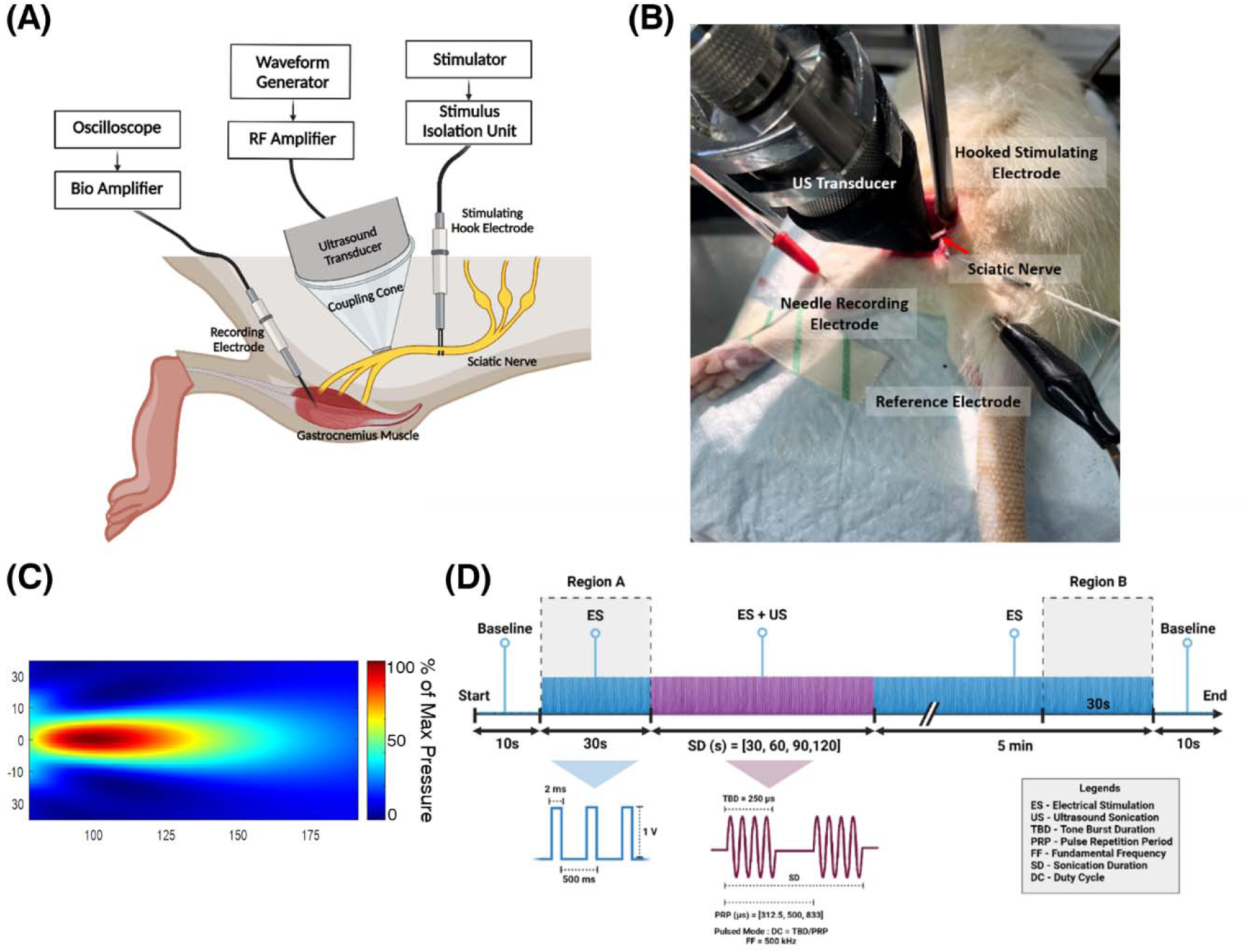
Experimental setup. (A) Neurophysiological recording schematic: The rat’s sciatic nerve receives electrical stimulation from a hooked stimulating electrode, and ultrasound sonication from an ultrasound transducer placed in close proximity of the nerve. A recording electrode inserted into the gastrocnemius muscle records changes in EMG. The electrical stimulation is driven via a stimulator connected to a stimulus isolation unit. The ultrasound transducer receives amplified continuous sine waves. (B) Animal experimental setup with the electrodes positioned in place for electrophysiological recording. (C) Acoustic field simulation, using Matlab toolbox K-wave, showing maximal pressure reached at a distance of 30 mm from the face of the transducer. (D) Stimulation procedure: Region A corresponds to 30 s of pure electrical stimulation. This was followed by concurrent ultrasound sonication for varied sonication periods. A 5 minutes interval of pure electrical stimulation was applied, with the last 30 s corresponding to Region B. A 10 s baseline recording initiated and concluded each stimulation session.

### 2.3. Ultrasound Sonication

A single-element unfocused ultrasound transducer (Mana Instruments E0525-SU) was connected to a 50-dB gain RF amplifier (Electronics & Innovation Ltd., Rochester, NY, USA), receiving electrical pulses from a waveform generator (SIGILENT-SDS 1025). A cone, of outer diameter 25 mm, coupled the transducer, of center frequency 500 kHz, with the sciatic nerve as it received ultrasonic pulses at a frequency of 500 kHz, number of cycles (CPP) = 125, and tone burst duration (TBD) = 250 µs. The cone, with a 2 mm inner diameter at the tip and a height of 30 mm, concentrated the ultrasound waves, precisely focusing the acoustic energy directly on the nerve. The sonication duration was varied between 30, 60, 90, and 120 seconds. The duty cycle of ultrasound waves was varied between 30%, 50%, and 80% by altering the burst period. SD and DC were modified, one at a time, to explore the offline impact of ultrasound on the sciatic nerve. Before the experiment, the ultrasound transducer coupled with the cone and filled with ultrasound gel, was submerged into a tank filled with degassed water. This allowed the measurement of the maximal intensity at the tip of the cone, representing the acoustic energy delivered to the sciatic nerve. Accurate measurements were conducted using a calibrated needle hydrophone (Onda HNR-0500), controlled by a 3D-motorized system that converts mechanical signals into electrical signals for processing. The spatial peak pulse average intensity (I_SPPA_)) was computed from the voltage measurements, averaged across the tone burst duration. Two intensities were considered, 1.4 W/cm² and 3.39 W/cm². The simulated pressure field showed the highest recorded pressure concentrated at a depth of 30 mm from the transducer’s surface (Figure 1C), producing a focal spot with a Full Width at Half Maximum (FWHM) of 4.5 mm.

### 2.4. Experimental and Targeting Procedure

The electrical and ultrasound stimulation protocol in each experiment had a maximum duration of 470 seconds, contingent upon the applied sonication duration (Figure 1D). To establish a baseline, a 10-second recording was conducted, at both the start and conclusion of each trial. Subsequently, a 30-second phase of exclusive electrical stimulation, denoted as Region A, was implemented. This was followed by ultrasound sonication lasting for the specified SD, concurrently administered with electrical stimulation. The DC was varied accordingly to examine the neuromodulatory effects of ultrasound on the sciatic nerve. Following that, a 5-minute interval of pure electrical stimulation ensued, with the last 30 seconds corresponding to Region B. This segment was designed to probe the prolonged effects of ultrasound sonication on the sciatic nerve.

### 2.5. Data Processing and Analysis

EMG recordings were obtained through a 3-1401 DAC (CED) and relayed back to Spike2 software. Subsequent signal processing and analysis were conducted offline using custom MATLAB scripts. Each rat underwent three trials, maintaining a consistent sonication duration while altering the DC in each trial. Calculations were performed at 30-second intervals, capturing 60 spikes (frequency = 2Hz) in each interval. For every spike in the EMG vector, both the peak-to-peak amplitude (V_pp_), indicative of the potential difference across the muscle, and the area under the curve (AUC), representing the maximum power per unit current circulation, were computed, excluding the stimulus artifact. Mean values were derived from each 60-spike interval. In each trial, the normalized percent average change in V_pp_ and AUC were calculated. This involved subtracting the average values in Region B from those in Region A, normalized over the values in Region A, effectively mitigating substantial inter-animal variances.

The cumulative energy exposure equation was used to account for both SD and DC when describing the energy of ultrasound waves on a unit area of tissue. This equation calculates the total energy E delivered to the tissue over a given sonication period, considering both the intensity of the ultrasound waves and the duration of exposure, where E = SD x DC x I_SPPA_. This energy term helps assess the potential effects of ultrasound on biological tissues.

### 1.1 Temperature Analysis

A K-type needle thermocouple (40 AWG/0.08 mm; Omega, Newark, NJ, USA) of 0.08 mm thickness was positioned in proximity between the sciatic nerve and the tip of the transducer cone, precisely at the targeting site and acoustic focus, for measuring temperature changes during ultrasound exposure. The thermocouple was connected to a USB-TC01 device (National Instruments, Austin, Texas), a channel temperature input device that receives the signal from the thermocouple wire featuring Instant-DAC technology for instant temperature readings. Temperature elevation resulting from ultrasound sonication, using the maximal parameters of 120 seconds sonication duration and 80% duty cycle, was quantified for sciatic nerves *in vivo*. Temperature measurements were recorded every second over a total duration of 160 seconds, with baseline measurements taken 20 seconds before and after ultrasound sonication.

### 2.6. Histology and Safety Analysis

Hematoxylin and Eosin (H&E) staining was performed to evaluate potential tissue damage resulting from ultrasound sonication, focusing on alterations in tissue morphology such as cellular swelling, necrosis, and inflammation. Upon concluding the experiment, the sciatic nerve was carefully extracted and immersed in 10% formaldehyde for a 24-hour fixation period. Subsequently, the axons were carefully teased out and prepared for H&E staining. Slide analysis was conducted using the uScopeMXII digital microscope at a magnification of 40x. 40x.

## 3. Results

### 3.1. Excitation vs. Suppression

We delivered low-intensity, 500 kHz ultrasound to the sciatic nerve and assessed its neuromodulatory effects in conjunction with electrical stimulation. Changes in electrically-evoked EMG of gastrocnemius muscle were measured following sciatic nerve exposure to ultrasound. The nerve exhibited varied neuromodulation responses, including enhanced electrical excitation, characterized by an increase in electrically induced EMG V_pp_, and suppression of electrically evoked excitation, marked by a decrease in electrically induced EMG V_pp_. For shorter SDs of 30s and 60s, the percentage average change in V_pp_ exhibited an increase across all applied DCs, peaking at 6.2% with an (SD, DC) combination of (30s, 80%). Conversely, longer SDs of 90s and 120s resulted in a decrease in the average change of V_pp_ for all applied DCs. The maximum suppression percentage of 5.1% was achieved with the (SD, DC) combination of (120s, 80%).

The response map in figure 2 demonstrates a bimodal EMG response, shifting from excitation to inhibition depending on the applied (SD, DC) combination. This transition from increased to suppressed EMG activity in response to ultrasound sonication becomes more pronounced with higher SD values. While increasing DC does enhance the degree of neuromodulation, it alone does not account for the shift in EMG activity following sonication. A two-way ANOVA showed a significant effect of SD on V_pp_ (p<0.001), however, no significant effect of DC was observed (p=0.751), suggesting that changes in DC alone do not significantly affect the percent average change in V_pp_. Interestingly, a significant interaction effect between SD and DC was significant (p=0.016), indicating that their combination has a different impact on V_pp_ that is not explained by either factor alone. Additionally, Tukey’s post-hoc analyses revealed significant differences between all levels of SD. Conversely, no significant differences were found between any of the DC levels, confirming that DC alone does not produce substantial changes in V_pp._

**Figure 2.**
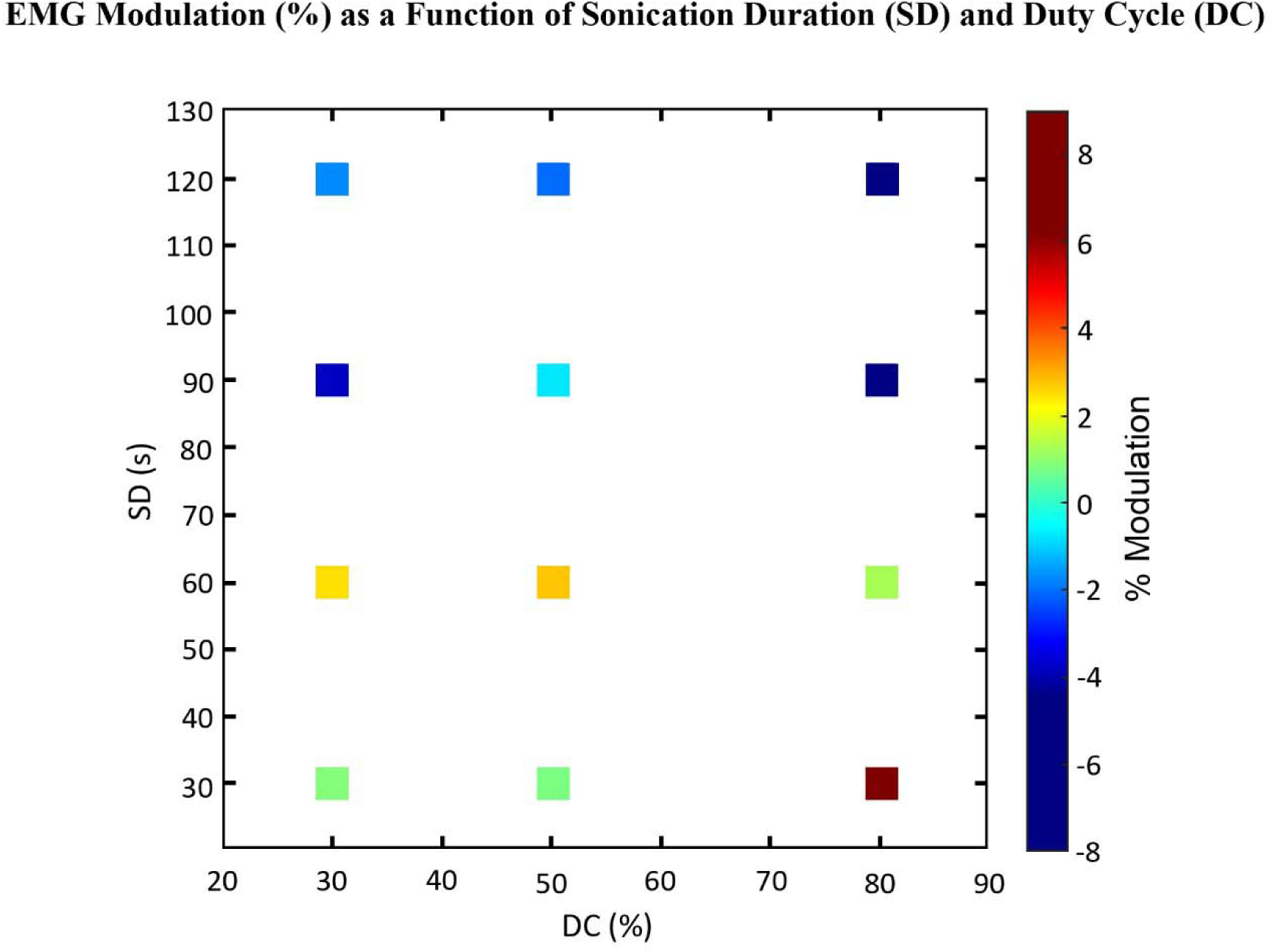
Response map showing the percent modulation based on percent average change in V_pp_ as a function of SD and DC. The spectrum shifts from stimulation of EMG activity at low SDs and DCs to suppression of EMG activity at higher SDs and DCs.

### 3.2. Effect of Acoustic Intensity on Nerve Activity

Acoustic intensity serves as a direct and immediate factor in ultrasound neuromodulation, driving neuromodulatory effects through mechanical forces and activation of mechanosensitive ion channels. The effect of varying acoustic intensity (I_SPPA_) on nerve activity was evaluated by measuring changes in EMG responses at two intensities: 1.4 W/cm² and 3.39 W/cm². Increasing the acoustic intensity significantly influenced both the percent average change in peak-to-peak EMG voltage V_pp_ and the area under the EMG response curve (AUC), demonstrating intensity-dependent neuromodulatory effects (Figure 3). At an intensity of 3.39 W/cm², the response was more pronounced, with a substantial increase in EMG excitation at shorter sonication durations and lower duty cycles, while longer sonication durations resulted in marked suppression of EMG activity. Specifically, for a 30s SD and 80% DC combination, V_pp_ increased by 137%, indicating enhanced excitation. In contrast, at the maximum sonication duration of 120s, the same intensity led to a 141.8% decrease in V_pp_,, illustrating a switch to inhibition (Figure 3A). Similarly, AUC measurements reflected this intensity-dependent modulation. With short sonication, AUC increased by 9.3% at 1.4 W/cm² and rose further to 64.1% at 3.39 W/cm² with extended sonication Figure 3. Response map showing the percent modulation based on percent average change in V_pp_ as a function of SD and DC. The spectrum shifts from stimulation of EMG activity at low SDs and DCs to suppression of EMG activity at higher SDs and DCs.

**Figure 3.**
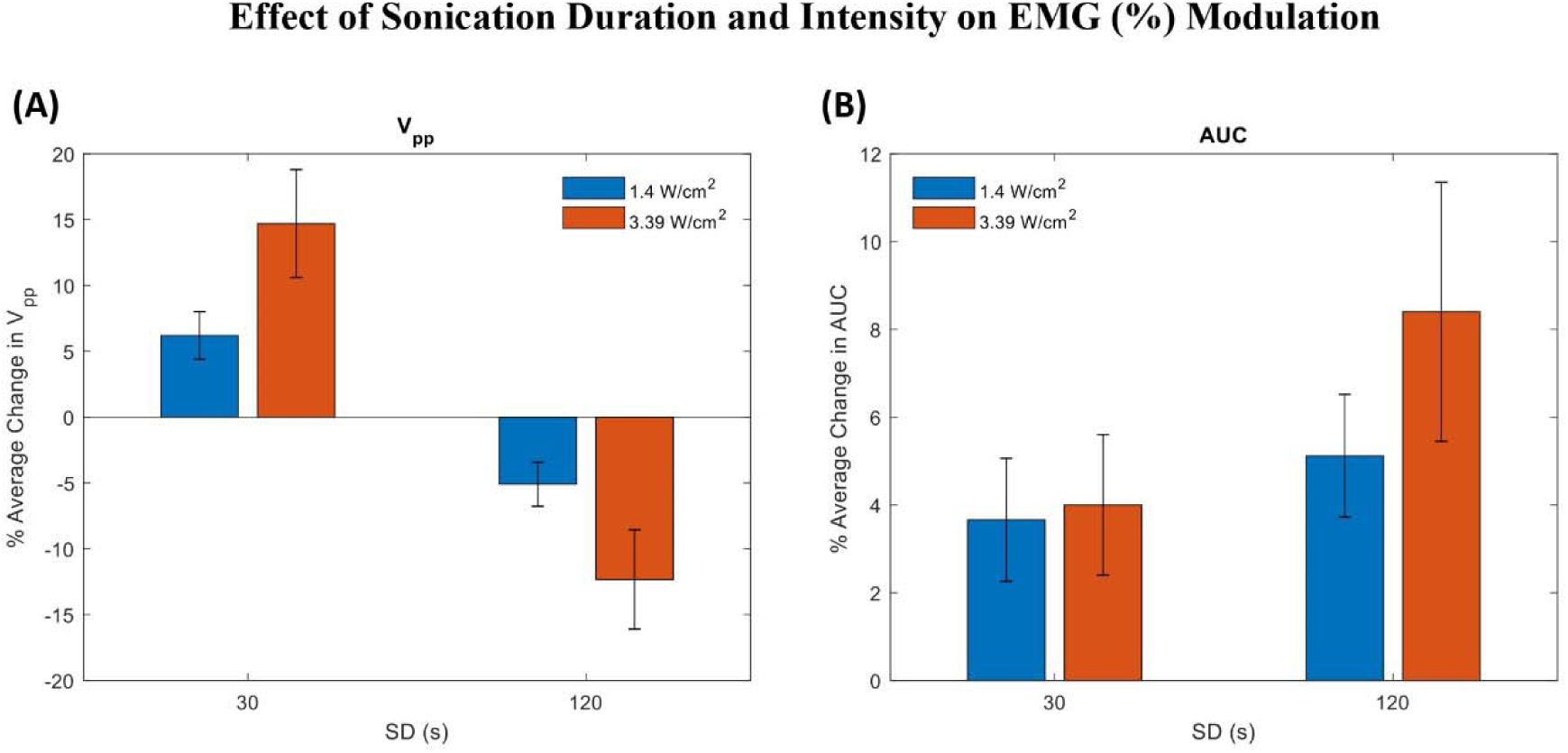
Percentage average change in (A) V_pp_ and (B) AUC as a function of I_SPPA_. Increasing the spatial-peak pulse-average intensity from 1.4 W/cm^2^ to 3.39 W/cm^2^ caused a significant change in the EMG activity at the two extreme sonication treatments with (SD, DC) of (30s, 80%) and (120s, 80%) respectively

(Figure 3B). This progression highlights the role of intensity in modulating the extent of neuromodulation, indicating that higher acoustic intensities facilitate a broader dynamic range of responses, from excitation to inhibition, based on the specific sonication parameters applied.

### 3.3. Prolonged Activity of Low-Intensity Low-Frequency Ultrasound

The modulation of EMG activity in the gastrocnemius muscle persisted for up to 5 minutes after low-intensity, low-frequency ultrasound was applied to the sciatic nerve, with a concurrent electrical stimulus. High cumulative energy levels resulted in an immediate decrease in EMG amplitude, with no further decline observed during the session (Figure 4A). A closer examination of individual muscle activity units in Regions A and B (Figure 4B) showed a 5.1% decrease in V_pp_. A more pronounced and sustained 28.7% decrease in EMG response was recorded (Figure 4C-D) when intensity was increased, further raising the energy level. Conversely, lower cumulative energy levels, at the same intensity, led to a 15.2% increase in EMG activity, maintained for 5 minutes post-sonication (Figure 4E-F). These results suggest that the delivered ultrasound dose can modulate and sustain the suppressive or excitatory response in muscle activity for an extended period.

**Figure 4.**
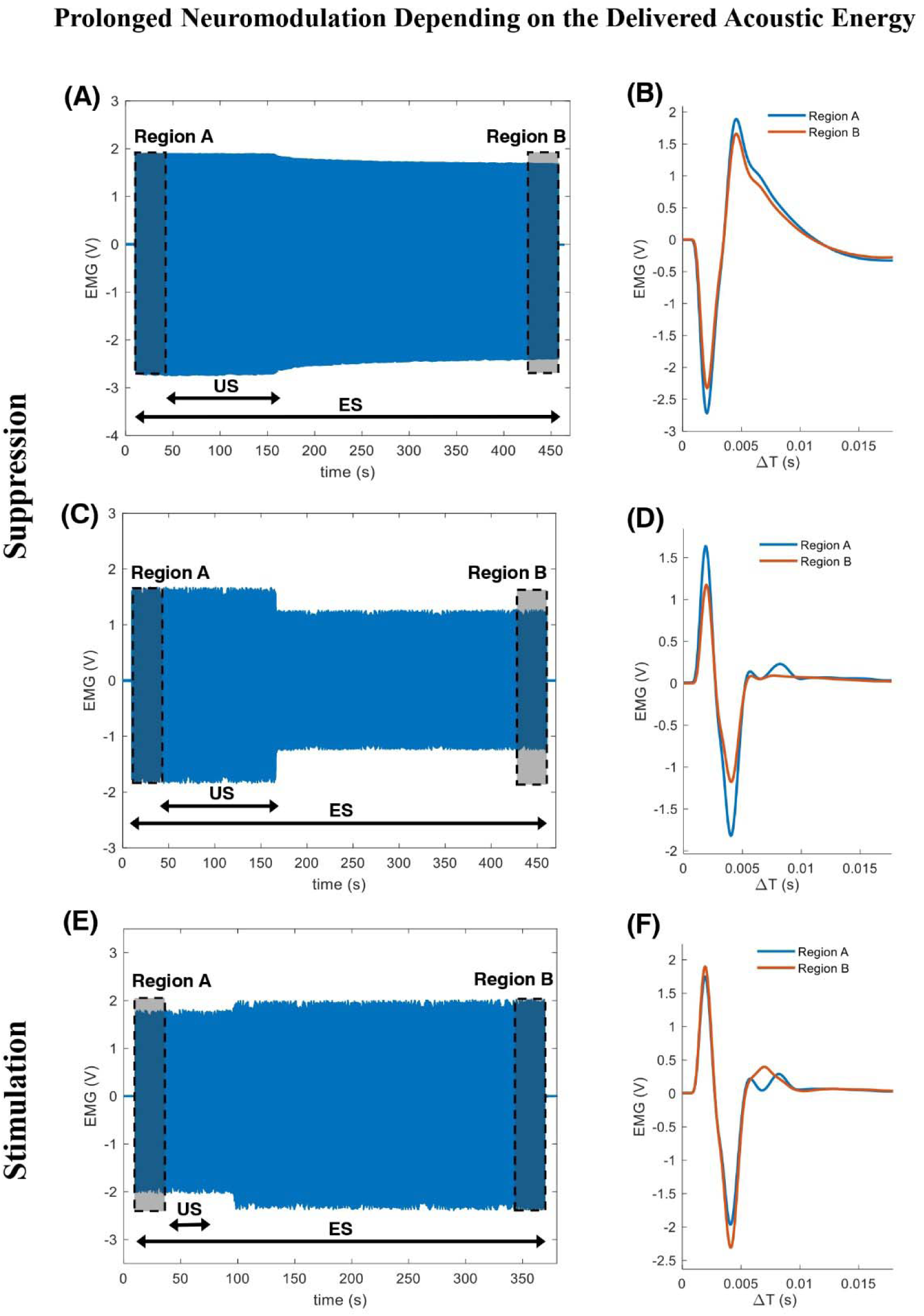
Sustained effect of ultrasound on EMG activity. (A) Prolonged suppression of EMG at high energy levels. (B) Single-unit muscle activity in Regions A and B shows a 5.1% drop in V_pp_. (C) Higher suppression of EMG activity was recorded when intensity was increased. (D) Zoomed view of a single EMG unit reveals a substantial 28.7% drop in V_pp_. (E) Prolonged excitation of EMG activity at lower energy levels. (F) Single-unit muscle activity in Regions A and B shows a 15.2% increase in V_pp_.

### 3.4. Effect of Ultrasound-Induced Energy on **V_pp_** and AUC

Each trial had its combination of SD and DC, thus a distinct cumulative energy-per-unit area. Our findings indicate that ultrasound dose significantly modulates electrically induced motor activity. Specifically, ultrasound can either enhance or diminish the effects of electrical stimulation (Figure 5A5A). Lower doses, below an ultrasound energy value of 40 J/cm²,², tend to amplify the response, whereas higher doses, above an ultrasound energy value of 67 J/cm², likely reduce it. At moderatemoderate ultrasound doses, however, individual variations in SD and DC become crucial factors, creating a transitional region where nerve activity varies between enhanced activation and suppression in response to ultrasound sonication.

**Figure 5.**
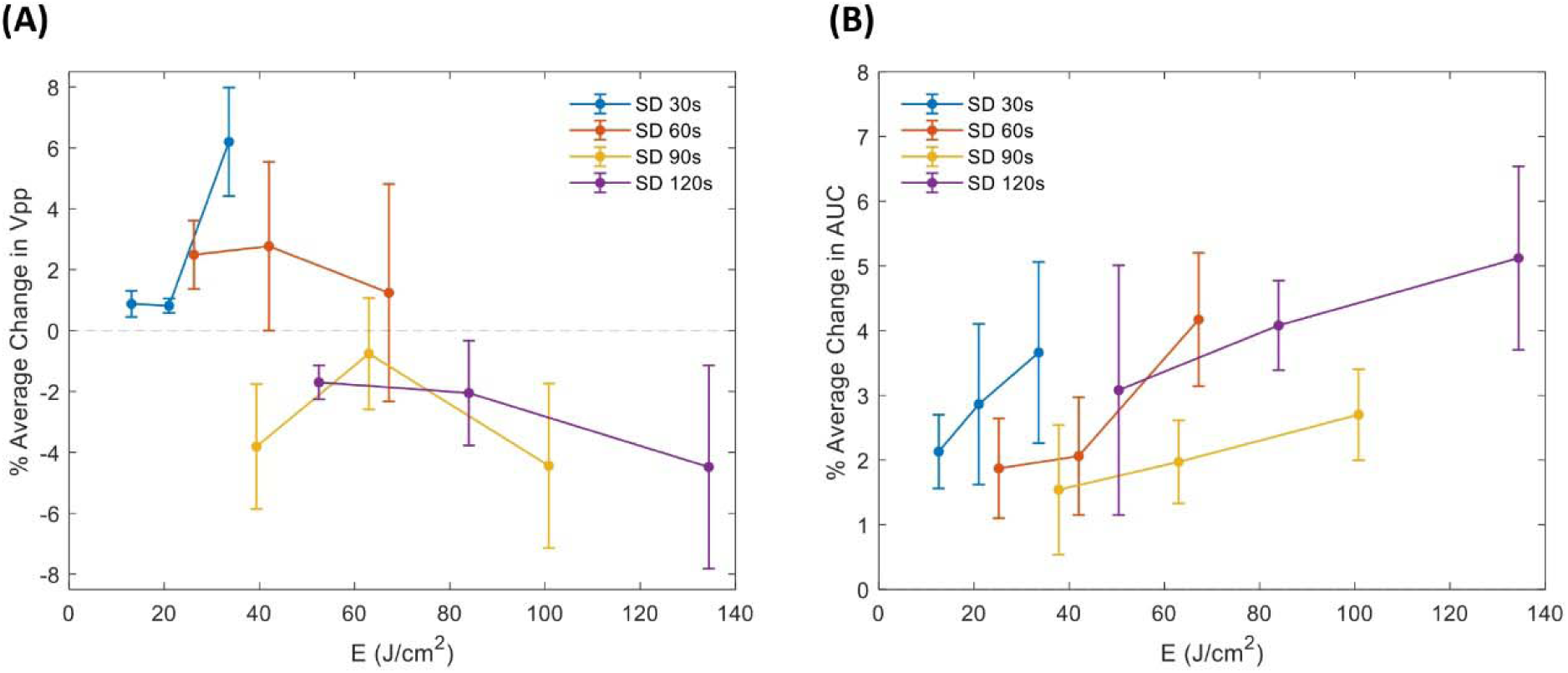
Ultrasound suppression and excitation of EMG activity based on ultrasound-induced cumulative energy exposure. (A) Percentage average change in V_pp_ as a function of energy dose. Energy term, represented with J_SPTA_ x SD, recorded an increase following an increase in SD and DC, shifting the EMG response from stimulation to suppression. (B) Percentage average change in AUC as a function of energy dose. Higher energy levels tend to produce greater changes in AUC across all sonication durations.

The EMG AUC was significantly affected depending on the ultrasound dose delivered. There was a general upward trend where the percentage average change in AUC increased following an increase in the cumulative energy delivered (Figure 5B). Higher energy levels tend to produce greater changes in AUC across all sonication durations, implementing wider EMG curves and suggesting that ultrasound may modulate the peak EMG response and the total activity within the response window. The highest (SD, DC) combination of (120s, 80%) recorded the highest AUC percentage increase of 5.1 %. The ‘on time’ of ultrasound during sonication affected the shape of the EMG waveforms, with more stretching as energy was increased.

### 3.5. Nerve Recovery **and**and Safety Assessment

A 15-minute break was implemented between each trial to facilitate muscle rest and the replenishment of neurotransmitters at the synapse level. The peak-to-peak amplitude in Region B of the preceding trial was compared with that in Region A of the subsequent trial, both conducted on the same rat. Notably, the absolute average change in peak-to-peak voltage and AUC registered non-zero values, indicating that the ultrasound effect diminished during the inter-trial break (Figure 6 A-B), and the nerve recovered to its initial state pre-sonication despite the delivered ultrasound energy during sonication.

**Figure 6.**
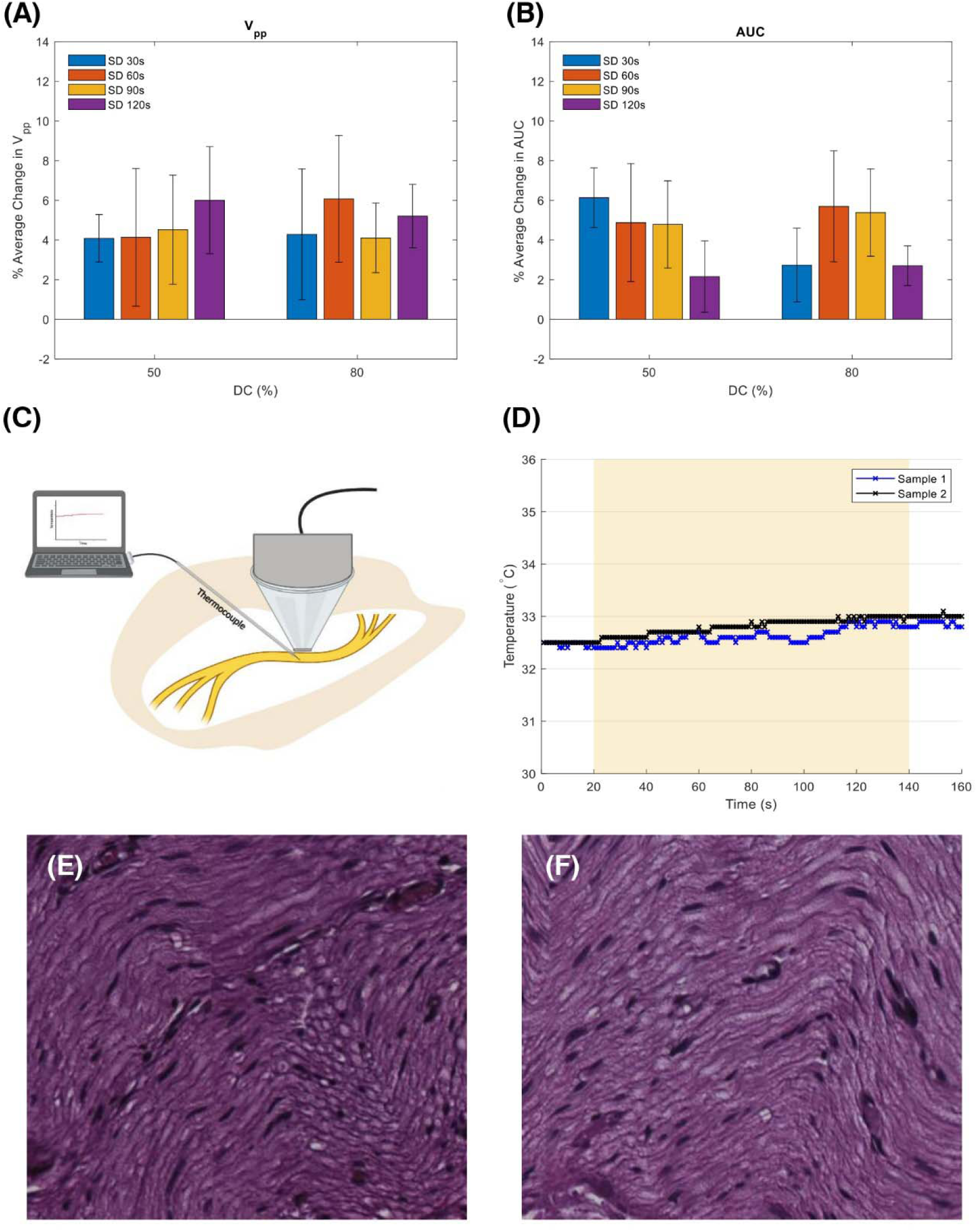
Reversibility and Safety of Ultrasound Neuromodulation. (A) Average percentage change in peak-to-peak voltage (V_pp_) and (B) area under the curve (AUC) as a function of sonication duration and duty cycle following a 15-minute nerve recovery period. (C) Temperature elevation (°C) measured with a K-type thermocouple placed between the ultrasound transducer and sciatic nerve. (D) Minimal temperature increase recorded during 120s of sonication at 80% DC in vivo for two sciatic nerve samples. (E) H&E staining of a sciatic nerve sample exposed to electrical stimulation only, and (F) staining of a sample exposed to both electrical stimulation and ultrasound sonication at the highest SD (120s) and DC (80%).

A thermal analysis was conducted to assess temperature changes resulting from ultrasound sonication of the sciatic nerve with the thermocouple placed at the targeting site and acoustic focus (Figure 6C). Minimal temperature variation was observed after applying ultrasound for 120 seconds at a DC of 80%. The two in vivo samples exhibited a maximum temperature increase of 0.34°C, shifting from 32.48°C ± 0.04°C to 32.7°C ± 0.16°C (Figure 6D). This gradual and minimal temperature elevation suggests the absence of tissue damage attributed to heat absorption. An intensity of 1.4 W/cm² corresponds to a low thermal index, implying a low probability of neuromodulation due to any induced thermal rise.

The safety of ultrasound application was evaluated through histological analysis using H&E staining. Examination of the H&E-treated images revealed no anatomical damage in the sciatic nerve tissue following ultrasound treatment (Figure 6 E-F). Under high magnification, there were no signs of inflammation or nerve degeneration.

## 4. Discussion

### 4.1. Energy-Based Ultrasound Neuromodulation

This study demonstrates a finely controlled, energy-dependent modulation of electrically evoked motor neuronal activity in the rat sciatic nerve achieved through low-intensity, low-frequency ultrasound. By systematically adjusting sonication duration and duty cycle, we identified a significant shift from excitatory to suppressive effects on EMG activity. This clear boundary between stimulation and inhibition based on cumulative energy exposure introduces a mechanistically robust framework for non-invasive peripheral neuromodulation, potentially offering a novel therapeutic approach with precise, controllable outcomes. Our findings underscore ultrasound’s capacity to affect sustained, bidirectional modulation of neural activity—either enhancing or inhibiting responses—without substantial increases in temperature, highlighting a primarily mechanical rather than thermal basis for neuromodulation. This contrasts with other approaches that rely on thermal or electrical mechanisms to influence neuronal behavior [20].

The cumulative energy exposure emerged as a pivotal factor in determining the type and extent of neuromodulation. At lower cumulative exposures (< 40 J/cm²), the data suggest that ultrasound facilitated electrically induced neural excitation. Shorter pulses can have higher peak power levels, even if the average power remains low. This high peak power can create stronger mechanical forces (such as acoustic radiation force) that are more effective at displacing or deforming nerve tissue, leading to excitation [21, 22]. This excitatory response may arise from mechanical activation of specific ion channels, likely mechanosensitive ones, which can alter membrane potentials and potentially enhance neurotransmitter release [23]. The fact that excitation occurred without heat buildup points to a primarily non-thermal, mechanotransductive mechanism, whereby acoustic forces directly interact with neuronal membranes or associated mechanosensitive structures, leading to heightened neuronal excitability [24–27].

Conversely, higher energy exposures (> 67 J/cm²) were associated with a marked suppression of EMG activity, which could stem from the increased mechanical load on neuronal membranes. This sustained acoustic radiation pressure and associated acoustic streaming might impede the natural flow of ions across the membrane by impacting membrane potential and ion channel conductance [28, 29]. Additionally, prolonged activation of mechanosensitive ion channels could lead to a degree of desensitization or receptor fatigue, thereby diminishing the nerve’s responsiveness to subsequent stimuli [30]. We propose that the delivered cumulative energy was converted into particle displacement, stretching ion channels at rates dependent on their composition. Indeed, ultrasound pressure waves can modulate transmembrane currents by influencing the kinetics of specific ion channels [31–33]. Yoo et al. demonstrated that inhibiting certain CNS ion channels reduces ultrasound response, with activation or inactivation affected by ultrasound-induced local particle displacement [34].

The interplay between excitation and suppression of electrically evoked EMG activity operates within a closed system highly dependent on the cumulative energy exposure. Our findings introduce a “dose-dependent switch” in neuromodulation, where specific energy thresholds enable a predictable shift from excitation to inhibition. This bimodal response to cumulative energy exposure, combined with the absence of significant thermal effects, positions ultrasound as a precise and adaptable modality for neuromodulation, expanding its potential as a non-invasive tool for therapeutic intervention.

A survey of existing literature [6, 7, 13, 20, 35–42] corroborates this bimodal trend, showing similar patterns of excitation at lower energy levels and suppression at higher energy levels across different neural models (Figure 7). For instance, studies in peripheral nerves and cortical regions have reported that low-energy ultrasound pulses facilitate neuronal excitability by engaging mechanosensitive ion channels, while higher energy levels result in suppression due to potential mechanical stress and receptor desensitization. These findings align closely with our experimental observations, further supporting the hypothesis that cumulative energy exposure plays a critical role in determining the neuromodulatory outcome of ultrasound stimulation. The variability in neuromodulation percentage for specific energy levels can be attributed to differences in experimental conditions across studies, such as tissue type, measurement techniques, or the targeted regions.

**Figure 7.**
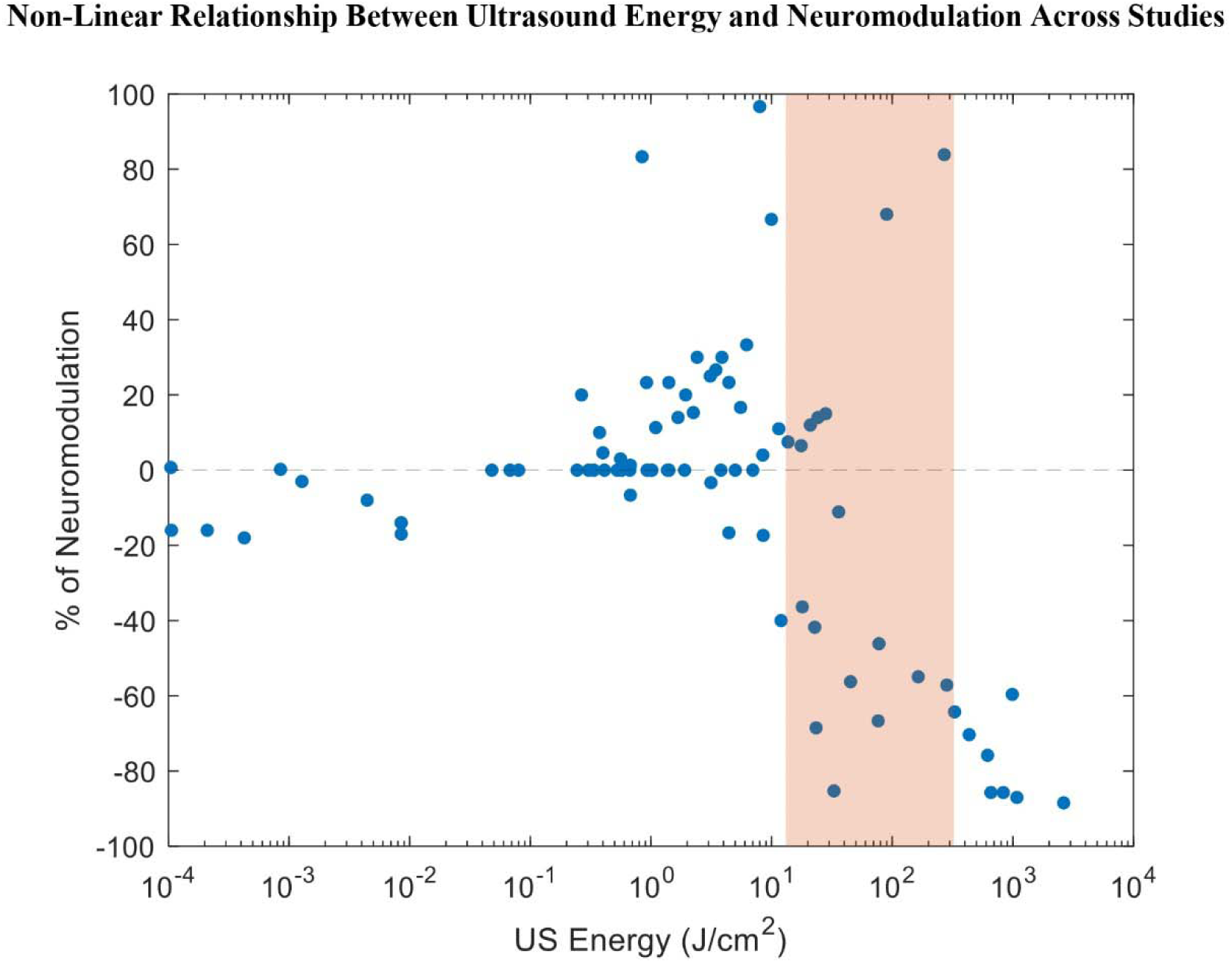
Neuromodulation Percentage as a Function of Ultrasound Energy (J/cm²) Across Studies. This plot displays the percentage of neuromodulation induced by ultrasound as a function of cumulative energy exposure, based on data from multiple published studies targeting both the central (CNS) and peripheral nervous systems (PNS). The data reveal a non-linear relationship, with variations in neuromodulatory effects at different energy levels. The shaded region highlights the energy range utilized in the current study, illustrating its positioning within the broader context of neuromodulation research.

### 4.2. Prolonged Effect of Ultrasound Neuromodulation

Low-intensity, low-frequency ultrasound applied to the rat sciatic nerve, combined with a consistent electrical stimulus, maintained muscle excitability or suppression for up to five minutes after cessation of sonication, with effects depending on the ultrasound dose. Following this, EMG activity gradually returned to baseline, indicating the dissipation of accumulated energy.

Ultrasound is known to activate cellular signaling pathways, inducing sustained changes in nerve excitability. Yoo et al. demonstrated that ten minutes of transcranial focused ultrasound on anesthetized rats resulted in sensory-evoked potential changes lasting up to 35 minutes [43], while Clennell et al. found that ultrasound enhances neuronal excitability by modulating ion channels, promoting synaptic plasticity, and increasing excitatory transmission in primary rat cortical neurons [44].

Studies on low-intensity focused ultrasound in non-human primates (NHPs) have shown that sonication of brain regions like the supplementary motor area and amygdala can alter connectivity patterns and functional coupling across regions [45]. This effect, observed in both NHPs and humans, suggests that ultrasound may modify neuronal responsiveness to other inputs rather than directly stimulating or inhibiting activity. The lasting effects of ultrasound may involve plastic changes such as long-term potentiation or depression, as seen in human prefrontal cortex stimulation, with no reported significant side effects. This effect, observed in both NHPs and humans, suggests that ultrasound may modify neuronal responsiveness to other inputs rather than directly stimulating or inhibiting activity. The lasting effects of ultrasound may involve plastic changes such as long-term potentiation or depression, as seen in human prefrontal cortex stimulation, with no reported significant side effects [46].

Further, experiments with leech mechanosensory neurons showed sustained membrane depolarization after sonication, supporting Dedola et al.’s hypothesis that ultrasound might cause prolonged modifications in ion channels, leading to increased neuronal responsiveness [47, 48]. These findings suggest a cumulative effect of ultrasound that potentially impacts neuron excitability and plasticity.

## 5. Conclusion

In this study, we investigated the effects of low-intensity, low-frequency ultrasound on rat sciatic nerve activity in vivo, focusing on variations in intensity, sonication duration, and duty cycle. Our results indicate that cumulative energy exposure, primarily influenced by SD and DC, is key to modulating motor neuron responses. Specifically, higher cumulative energy levels suppressed EMG activity, while lower levels led to excitation, suggesting an energy-based mechanism for ultrasound neuromodulation.

These effects occurred without cavitation or thermal changes, implying a mechanical rather than thermal mode of action. Additionally, no detectable nerve damage was observed, confirming a safe range for prolonged activation or suppression of neural activity. Given our findings, we hypothesize that ultrasound could potentially act as a proxy for endogenous electrical stimulation, modulating naturally driven motor activity in a non-invasive manner. This opens possibilities for further research into ultrasound’s potential to replicate the body’s natural neural control mechanisms. Future studies should aim to clarify the exact mechanisms and optimize ultrasound parameters for precise neural modulation.

